# Cryopreservation of human cancers conserves tumour heterogeneity for single-cell multi-omics analysis

**DOI:** 10.1101/2020.06.04.135277

**Authors:** Sunny Z. Wu, Daniel L. Roden, Ghamdan Al-Eryani, Nenad Bartonicek, Kate Harvey, Aurélie S. Cazet, Chia-Ling Chan, Simon Junankar, Mun N. Hui, Ewan A. Millar, Julia Beretov, Lisa Horvath, Anthony M. Joshua, Phillip Stricker, James S. Wilmott, Camelia Quek, Georgina V. Long, Richard A. Scolyer, Bertrand Z. Yeung, Davendra Segara, Cindy Mak, Sanjay Warrier, Joseph E. Powell, Sandra O’Toole, Elgene Lim, Alexander Swarbrick

## Abstract

**Background:** High throughput single-cell RNA sequencing (scRNA-Seq) has emerged as a powerful tool for exploring cellular heterogeneity amongst complex human cancers. scRNA-Seq studies using fresh human surgical tissue is logistically difficult, precludes histopathological triage of samples and limits the ability to perform batch processing. This hinderance can often introduce technical biases when integrating patient datasets and increase experimental costs. Although tissue preservation methods have been previously explored to address such issues, it is yet to be examined on complex human tissues, such as solid cancers, and on high throughput scRNA-Seq platforms.

**Results:** We show that the viable cryopreservation of human cancers provides high quality single-cell transcriptomes using the Chromium 10X platform. We sequenced a total of ∼120,000 cells from fresh and cryopreserved replicates across three breast cancers, two prostate cancers and a cutaneous melanoma. Importantly, tumour heterogeneity identified from fresh tissues was largely conserved in cryopreserved replicates. We show that sequencing of single cells prepared from cryopreserved tissue fragments or from cryopreserved cell suspensions is comparable to sequenced cells prepared from fresh tissue, with cryopreserved cell suspensions displaying higher correlations with fresh tissue in gene expression. We then show that cryopreservation had minimal impacts on results of downstream analyses such as biological pathway enrichment. Further, we demonstrate the advantage of cryopreserving whole-cells for immunophenotyping methods such as CITE-Seq, which is impossible using other preservation methods such as single nuclei-sequencing.

**Conclusions:** Our study guides new experimental designs for tissue biobanking for future clinical single-cell RNA sequencing studies.

## Background

The tumour microenvironment (TME) is composed of neoplastic cells, parenchymal and immune cells that interact to shape cancer progression and therapeutic response [1]. Advances in high-throughput single-cell RNA sequencing (scRNA-seq) technologies have rapidly developed in recent years, providing a powerful platform to resolve the aetiology of the TME in solid cancers. Performing scRNA-seq on clinical samples remains logistically and technically challenging mainly due to transport of patient tissue from operation rooms to laboratories for processing, which are often complicated by short notices and core-facility access hours. The need to process fresh tissue specimens at the time of tissue availability, often as a single specimen, often introduces large experimental costs and confounding batch effects upon studies with large numbers of patients and prevents the selection and triage of cases for analysis based on histopathological analysis.

Several approaches have been developed to address such issues. Madissoon *et al.* benchmarked short-term cold preservation of tissue prior to scRNA-Seq, which showed little impact on transcriptome integrity within the first 24 hours [2]. Despite this, such short-term storage periods still limit the ability to perform simultaneous sample processing. Cell type specific transcriptional changes have been shown to emerge after longer cold preservation periods (>24 hours), particularly affecting immune subpopulations in normal tissues [2].Cold preservation is yet to be evaluated for complex tissues such as solid tumours, which possess distinct features in tissue viability. Factors including tissue necrosis, hypoxia and therapeutic treatments often result in poor viability of cells in solid tumour tissues. Cell fixation methods using agents such as methanol can be applied to overcome barriers of cold preservation. However, these methods are not always practical with solid cancers which require lengthy dissociation protocols, and also preclude certain downstream procedures such as antibody staining or cell culture [3, 4]. Although sequencing of nuclei from snap frozen tissue can be applied to avoid dissociation methods, this approach is not compatible with powerful cell surface immunophenotyping methods with DNA-barcoded antibodies such as CITE-Seq [5]. It also does not permit the selection of cell subsets of interest or the removal of low-quality cells prior to capture. Guillaumet-Adkins *et al.* showed that the cryopreservation of whole-cells and tissues can be used to conserve transcriptional profiles from experimental systems such as human cell lines and mouse tissues [6]. These models represent fairly homogeneous systems and it is unclear whether the highly heterogeneous nature of the TME is also conserved following cryopreservation. In addition, this study benchmarked tissue cryopreservation using low-throughput plate-based scRNA-seq technology [6], where highly viable cells are selected by FACS for immediate lysis and mRNA hybridisation [7]. It is yet to be determined if cryopreservation can be applied to more recent high throughput scRNA-Seq platforms such as the Chromium 10X platform. These platforms are fundamentally different to FACS-based scRNA-Seq methods, as single-cells are captured through droplet-based microfluidics, where viability selection is not simultaneously performed.

In this study, we aimed to examine the effect of cryopreserving dissociated cells and solid tissues prior to scRNA-Seq on the 10X Chromium platform. We tested this across three common cancer types: breast, prostate and melanoma. Following cryopreservation, we demonstrated a strong conservation of the heterogeneous neoplastic, parenchymal and immune subpopulations. We show that scRNA-Seq results of cells from cryopreserved solid tissue and from cryopreserved dissociated cell suspensions are comparable to those from cells prepared from fresh tissue, with minimal impact on downstream analysis methods. Lastly, we show that cryopreserving whole-cells allows for powerful immunophenotyping methods such as CITE-Seq, which is not possible using nuclei-based sequencing methods. Our findings allow a simple biobanking protocol to process patient samples, significantly decreasing technical variation among larger patient cohorts and serial time-points analyses. Our biobanking protocol unlocks patient cohorts previously collected in such a manner, and serves as a guide for the sample collection in future clinical scRNA-Seq studies.

## Results and Discussion

### Cryopreservation allows for robust conservation of cellular heterogeneity in human breast cancers

The preservation of cellular heterogeneity is an important factor for analysing solid cancers. We first investigated this in primary human breast cancers collected from three patients (Supplementary Table 1). To minimise spatial biases from regional sampling, fresh surgical specimens were initially cut in to 1-2 mm^3^ pieces and thoroughly mixed. One third of the mix was immediately cryopreserved at -80°C (designated as the cryopreserved tissue - CT) and the remaining mix was dissociated into a single-cell suspension using a commercial kit-based method (See Methods). A fraction of this cell suspension was immediately cryopreserved at -80°C (designated as the cryopreserved cell suspension - CCS) and the remaining of this cell suspension was processed immediately for scRNA-Seq using the Chromium 10X platform (designated as fresh tissue - FT). After storage of the cryopreserved samples, both CT and CCS, at -80°C for about one week, they were stored in liquid nitrogen at -196°C for up to five weeks to mimic standard tissue biobanking procedures. Following cryopreservation, CT and CCS samples were thawed and processed for scRNA-Seq in the same manner as the FT sample. For cryopreserved replicates, we spiked in the mouse NIH3T3 fibroblast cell line as a positive control (∼2%) for the scRNA-Seq experimental workflow. In total, we sequenced 23,805, 29,865 and 24,250 cells from breast cancer patients 1-3, (assigned as BC-P1, BC-P2 and BC-P3), respectively.

A detailed comparison was performed between samples processed as FT, CCS or CT (Fig. 1a). We performed batch correction and integration of all matched fresh and cryopreserved replicates using the anchoring based method in Seurat v3 (Fig. 1b) [8]. This revealed consistent ‘mixability’ across the three conditions, where a strong overlap was also observed in Uniform Manifold Approximation and Projection (UMAP) space. This was also observed in the non-batch corrected data (Fig. S1a), reflecting good technical replicates on the 10X Chromium platform. To account for variation in cell-type proportions, all matched conditions were down sampled to the lowest replicate cell number to examine the composition of cells in each cluster (Fig. 1c). Only three clusters across all three datasets were not comprised of cells from all three conditions (Fig. 1c). These differential clusters were all detected in the BC-P2 dataset, including clusters c11 (737 cells), c18 (191 cells) and c23 (27 cells). Clusters c11 and c18 were only detected in the FT sample and resembled cell doublets captured from a varying number of cells sequenced per replicate, which ultimately contributes to a differences in the expected doublet rate. These clusters showed characteristics of cell doublets, including the expression of markers from multiple cell lineages such as *EPCAM, PTPRC, PECAM1* and *PDGFRB* (Fig. S1b). Cluster c23 was comprised of smaller cell numbers, and may be a result of sampling rarer cell types, rather than from the cryopreservation process. To our surprise, the mouse NIH3T3 fibroblast spike-ins could also be detected in all cryopreserved replicates following the mapping of reads to the human GRCh38 reference genome alone (c19 in BC-P1, c17 in BC-P2 and c14 in BC-P3). These were confirmed as mouse cells by re-mapping reads to both human and mouse reference genomes, suggesting that mouse reads were assigned to their human orthologs when mapping to a single reference genome using CellRanger. NIH3T3 fibroblast spike-ins captured from different cryopreserved replicates and independent experiments mixed well (Fig. S1c), indicating high reproducibility on the 10X Genomics platform. As expected, NIH3T3 fibroblasts highly expressed markers *Dlk1, Acta2, Vim, Actg1, Col1a1* and *Col1a2* (Fig. S1d).

**Figure 1.**
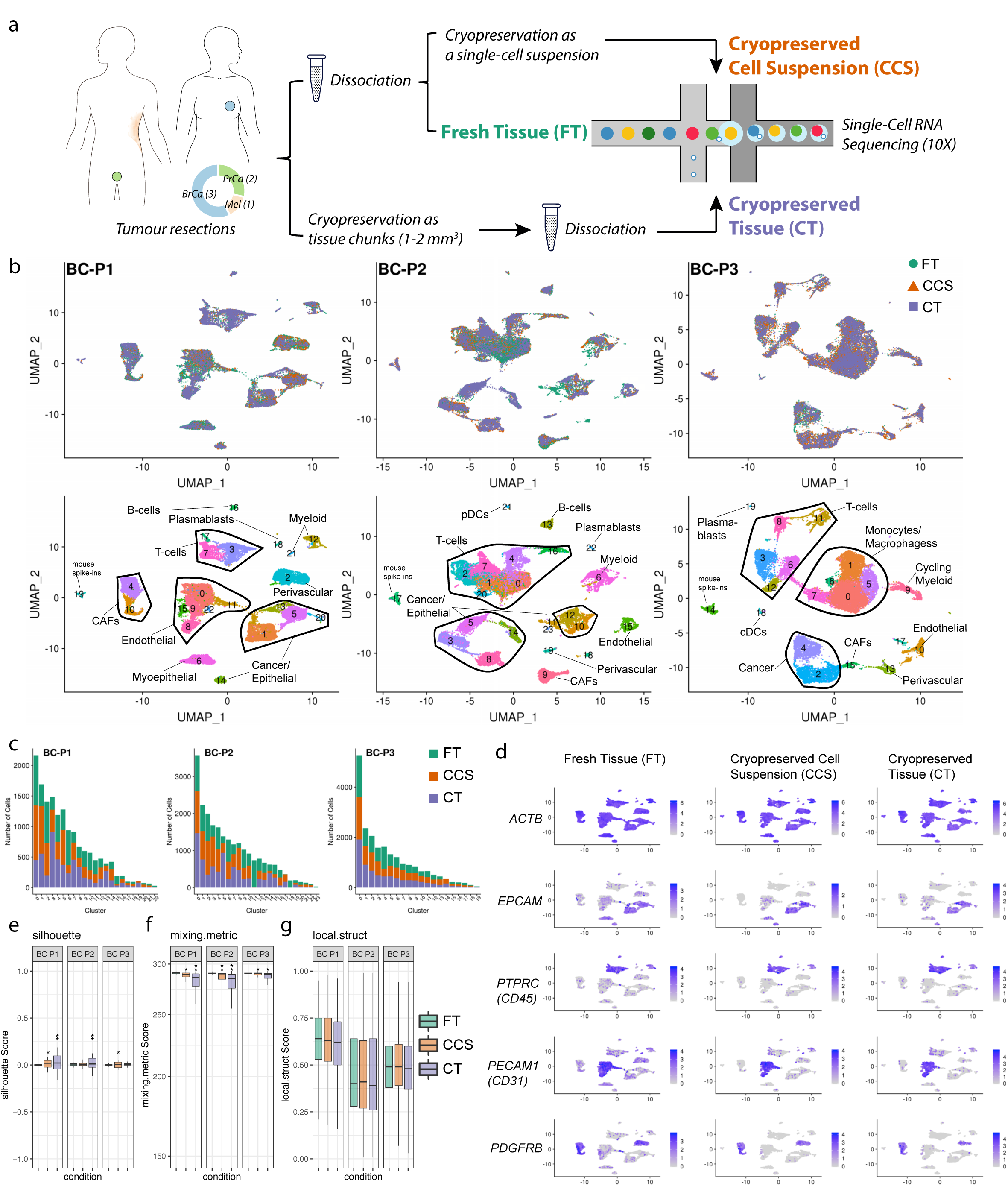
Cryopreservation allows for robust cell-type detection in clinical breast cancer samples. **a**, Experimental workflow. **b**, UMAP visualisation of 23,803, 29,828 and 24,250 cells sequenced across dissociated fresh tissue (FT; green), dissociated cryopreserved cell suspensions (CCS; orange) and solid cryopreserved tissue (CT; purple) replicates from three primary breast cancer cases (BC-P1, BC-P2 and BC-P3). UMAPs are coloured by cryopreserved replicate (top) and by cluster ID (bottom) with cell types annotations overlayed. Matched replicates were integrated using the Seurat v3 method. **c**, Number of cells detected per cluster. Cells were down sampled to the lowest replicate size. **d**, Featureplot visualisations of gene expression from BC-P1 fresh and cryopreserved replicates, showing the conservation of the housekeeping gene *ACTB* and heterogeneous cancer/epithelial (*EPCAM*), immune (*PTPRC/*CD45), endothelial (*PECAM1*/CD31) and fibroblast/perivascular (*PDGFRB*) clusters. **e-g**, Distribution of silhouette scores (range -1 to +1) **(e)**, mixing metric **(f)**, and local structure metrics **(g)** of clustering following cryopreservation. Samples were down sampled by replicate and cluster sizes and compared to the respective FT samples. Cell comparisons were performed across down sampled FT-1 vs FT-2 cells (positive control), FT vs CCS cells and FT vs CT cells. Stars represent standard deviations; **(e)** silhouette scores s.d. 0.02 - 0.05* and s.d. > 0.05**; **(f)** mixing metrics s.d. 2 - 10* and s.d. > 10**; **(g)** local structure metrics s.d. > 0.05*.

From investigating the expression of canonical cell type markers, we identified a strong preservation of major cell lineages in cryopreserved replicates (Fig. 1d). As observed in the representative case BC-P1 (Fig. 1d), we identified a strong conservation of the housekeeping gene *ACTB*, cancer/epithelial cells (*EPCAM;* clusters c1, c5, c13, c20 and c14), myoepithelial cells (*KRT14;* c6), T-cells (*CD3D;* c3, c7 and c17), B-cells (*MS4A1;* c16), plasmablasts (*JCHAIN*; c18), myeloid cells (*CD68;* c12 and c21), endothelial (*PECAM1;* c0, c8, c9, c11, c15 and c22), perivascular cells (*PDGFRB;* c2) and cancer-associated fibroblasts (CAFs; *PDGFRA;* c4 and c10) (Fig. 1d; Fig. S2a; Supplementary Table 2). Similar trends in the preservation of the TME was observed in all three breast cancer cases (Fig. S1b; Fig. S2b-c; Supplementary Table 2). In summary, cryopreservation of human breast cancers as either solid tissue or single cell suspension maintains the heterogeneity of major cell lineages detected from processing fresh tissue.

### Cryopreserved replicates resemble good technical replicates with the fresh tissue data

Although visual inspection of the dimensional reduction UMAP plots indicated good mixability and minimal technical variation emerging from cryopreservation, we applied several metrics adopted from Stuart *et al.* to quantitatively measure the impact on downstream clustering [8]. We examined silhouette coefficient scores, mixing metric and local structure metric to measure the robustness of cryopreservation to reflect good technical replicates with the FT. As described in the previous section, we performed stratified down sampling of cells to account for differences emerging from total number of cells sequenced. We compared cells from FT against cells from matched cryopreserved replicates independently in the following comparison conditions: FT vs CCS and FT vs CT. As a positive control, we compared two sets of FT cells down sampled from the same dataset to reflect perfect technical replicates (FT-1 vs FT-2).

Silhouette coefficient scores, which range from -1 to +1, measure how similar a cell is to cells from its own cluster in dimensional reduction space. We applied this to measure the mixability of the cryopreserved replicates, where scores closer to 0 indicate a higher mixability between replicates irrespective of cryopreservation condition. As expected from our positive control comparisons (FT-1 vs FT-2), this yielded average silhouette scores close to 0 for all three breast cancer cases (Fig. 1e). In general, we observed values close to 0 for all cryopreserved replicate comparisons, with no silhouette scores outside of the - 0.25 to 0.25 range (Fig. 1e). Minor variations, as indicated through increased standard deviations, were observed in the CCS replicates of two cases: BC-P1 and BC-P3 (Supplementary Table 3; Fig. 1e). Similarly, increased standard deviations were observed when comparing CT replicates in two cases: BC-P1 and BC-P2 (Supplementary Table 3; Fig. 1e).

We next applied the mixing metric to assess how well cryopreserved replicates ‘mixed’ with the FT data after integration (Fig. 1f). The mixing metric examines the distribution of replicates in a cell’s neighbourhood (k = 5 and k.max = 300), where values closer to 300 resemble a high ‘mixability’ (Fig. 1f) [8]. Overall, very high mixing metric scores were observed across the comparison conditions from all three breast cancer cases; however, slightly lower values and higher standard deviations were consistently detected in cells cryopreserved as CT compared to CCS (Supplementary Table 3; Fig. 1f). Finally, we assessed how local cell clusters (k = 100) detected in individual replicates were preserved upon data integration using the local structure metric [8]. In all three cases, this revealed no major differences in the standard deviations from our positive FT control comparisons and the cryopreserved replicates (Supplementary Table 3; Fig. 1g), indicating that the clusters identified in individual replicates were largely consistent upon integration with the FT data. Overall, we conclude that cryopreservation can yield good quality technical replicates. Only minor variation in clustering, as determined by Silhouette coefficients and mixing metrics, arise from processing as dissociated CCS and solid CT, with the latter resulting in slightly more variable data.

### Cryopreservation yields high quality data in prostate cancers and a metastatic melanoma

Tissue architectures differ across cancer sites and metastatic lesions. To assess the impact of cryopreservation across different tissue sites, we repeated our benchmarking on primary prostate cancer tissue collected from two patients (PC-P1 and PC-P2), and metastatic melanoma tissue collected from one patient (M-P1). For the metastatic melanoma sample, we benchmarked cell suspensions cryopreserved immediately (CCS sample) as well as after overnight storage of the tissue at 4°C in media (designated as cryopreserved overnight - CO). The CO replicate mimics conventional biobanking procedures where tissue is collected from late patient procedures, stored at 4°C and processed the following day. In total, we sequenced 18,333, 18,327 and 21,363 cells from PC-P1, PC-P2 and M-P1, respectively (Fig. 2a). Here, we identified that the CCS replicate from PC-P2 resulted in low cell number (less than 400) and was excluded from subsequent comparisons. Similar to the breast cancer data, comparisons of the non-batch corrected data revealed a good mixture of cells from all conditions, reflecting that of good technical replicates (Fig. S1a). Batch correction and data integration revealed consistent mixability across the three conditions in UMAP space (Fig. 2a-b; Fig. S1e). Only one very small cluster in PC-P1 (c20 – 64 cells) was not comprised of cells from all three conditions (Fig. 2c), and is, again, likely a result of cell sampling rather than cryopreservation. All clusters detected in M-P1 were comprised of cells from all conditions (Fig. 2c). Similar to our benchmarking in breast cancers, we observed a strong conservation of the housekeeping gene *ACTB* and markers for cancer clusters (*EPCAM* in prostate and *MITF* and melanoma), immune subsets (*PTPRC*), endothelial cells (*PECAM1*/CD31) and fibroblast/perivascular (*PDGFRB*) cells in prostate cancers and the metastatic melanoma (Fig. 2d-e; Fig. S2d-e). Upon examining clustering metrics, we found similar trends with slightly higher variation in silhouette scores and mixing metrics emerging from cells cryopreserved as CT compared to CCS (Fig. 2f-g; Supplementary Table 3). For the melanoma comparisons, the CO replicate exhibited a higher variation in silhouette scores and mixing metric compared to CCS, indicating potential transcriptional artefacts arising from overnight cold preservation prior to cryopreservation (Fig. 2f-g; Supplementary Table 3). No major differences were observed in the local structure metric of both prostate and melanoma cases (Fig. 2h), indicating that clustering neighbourhoods in individual replicates were consistently detected upon integration with the FT data. Taken together, our benchmarking across multiple tissue sites indicates that cryopreservation preserves the cellular heterogeneity of the TME and acts as good quality technical replicates.

**Figure 2.**
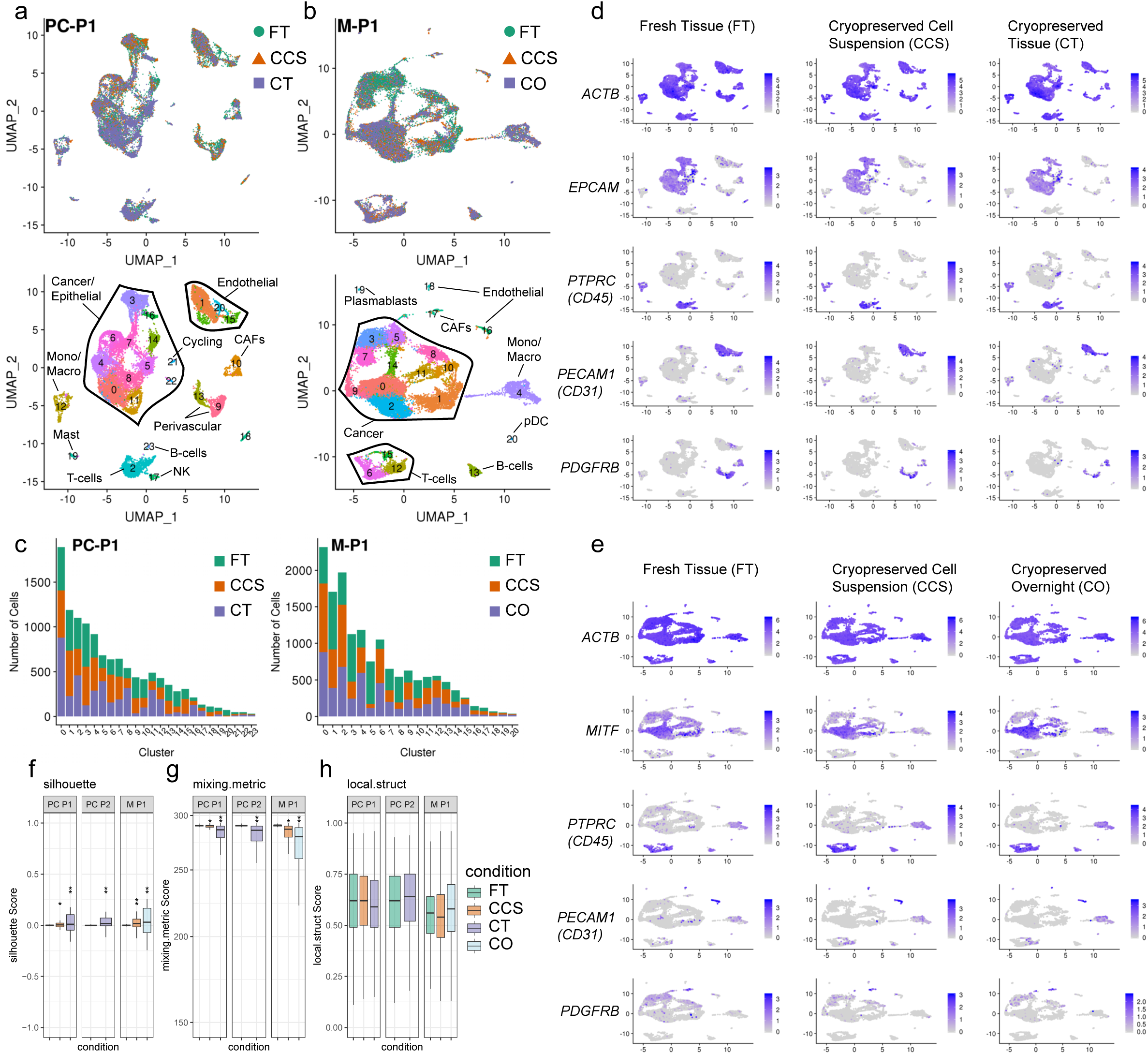
Cryopreservation allows for robust cell-type detection in clinical prostate cancer and melanoma samples. **a**, UMAP visualisation of 18,331 cells sequenced across FT (green), CCS (orange), and CT (purple) from primary prostate cancer case PC-P1. UMAPs are coloured by cryopreserved replicate (top) and by cluster ID (bottom) with cell types annotations overlayed. Matched replicates were integrated using the Seurat v3 method. **b**, UMAP visualisation as in **(a)** of 21,361 cells sequenced across FT (green), CCS (orange), and cryopreserved overnight (CO; purple) replicates from metastatic melanoma case M-P1. **c**, Number of cells detected per cluster from PC-P1 and M-P1, highlighting the conservation of clusters detected in the FT samples following cryopreservation. Cells were down sampled to the lowest replicate size. **d-e**, Featureplot visualisations of gene expression in prostate cancer **(d)** and melanoma **(e)** showing the conservation of the housekeeping gene *ACTB* and heterogeneous cancer/epithelial (*EPCAM* in **d** or *MITF* in **e**), immune (*PTPRC/*CD45), endothelial (*PECAM1*/CD31) and fibroblast/perivascular (*PDGFRB*) clusters following cryopreservation as FT, CCS and CT or CO. **f-h**, Distribution of silhouette scores **(f)**, mixing metric **(g)**, and local structure metrics **(h)** of clustering following cryopreservation as analysed in Fig. 1e-g. Stars represent standard deviations; **(f)** silhouette scores s.d. 0.02 - 0.05* and s.d. > 0.05**; **(g)** mixing metrics s.d. 2 - 10* and s.d. > 10**; **(h)** local structure metrics s.d. > 0.05*.

### Tumour cryopreservation maintains the integrity and complexity of single-cell transcriptomes

We next investigated whether gene expression and transcriptome integrity were affected through the cryopreservation process. We first examined the number of genes and unique molecular identifiers (UMIs) detected per cell across cryopreserved replicates. For this comparison, libraries were first down sampled by the number of mapped sequencing reads to account for differences emerging from varying sequencing depths. This revealed that an average of 1,809, 1,842 and 1,694 genes and 6,149, 6,525 and 5,851 UMIs per cells were detected across all FT, CCS and CT replicates, respectively (Fig. 3a-b). Within matched cases, only cryopreserved cell suspension replicates from M-P1 (from both CCS and CO) yielded a lower average number of genes and UMIs per cell compared to the FT (Fig. 3a-b). Similarly, only one CT replicate (BC-P1) had a significantly lower number of genes and UMIs detected per cell compared to the FT (Fig. 3a-b). Although this was not observed across multiple cases, a lower detection rate from CT may reflect a minor impact on transcript abundance and quality from the cryopreservation process. In addition, cell type and cell size can be an important factor determining transcript abundance. To determine that these subtle changes were not due to differences in cell abundance across cryopreserved replicates, we confirmed that these changes were also present at the cluster level (Fig. S3). For example, although cancer cells (clusters c1, c5 and c14 in BC-P1) generally hold more transcripts compared to T-cells (clusters c3, c7 and c17 in BC-P1), less genes and UMIs were also found in these respective cell types captured in CT replicate, as per the bulk comparisons (Fig. S3a).

**Figure 3.**
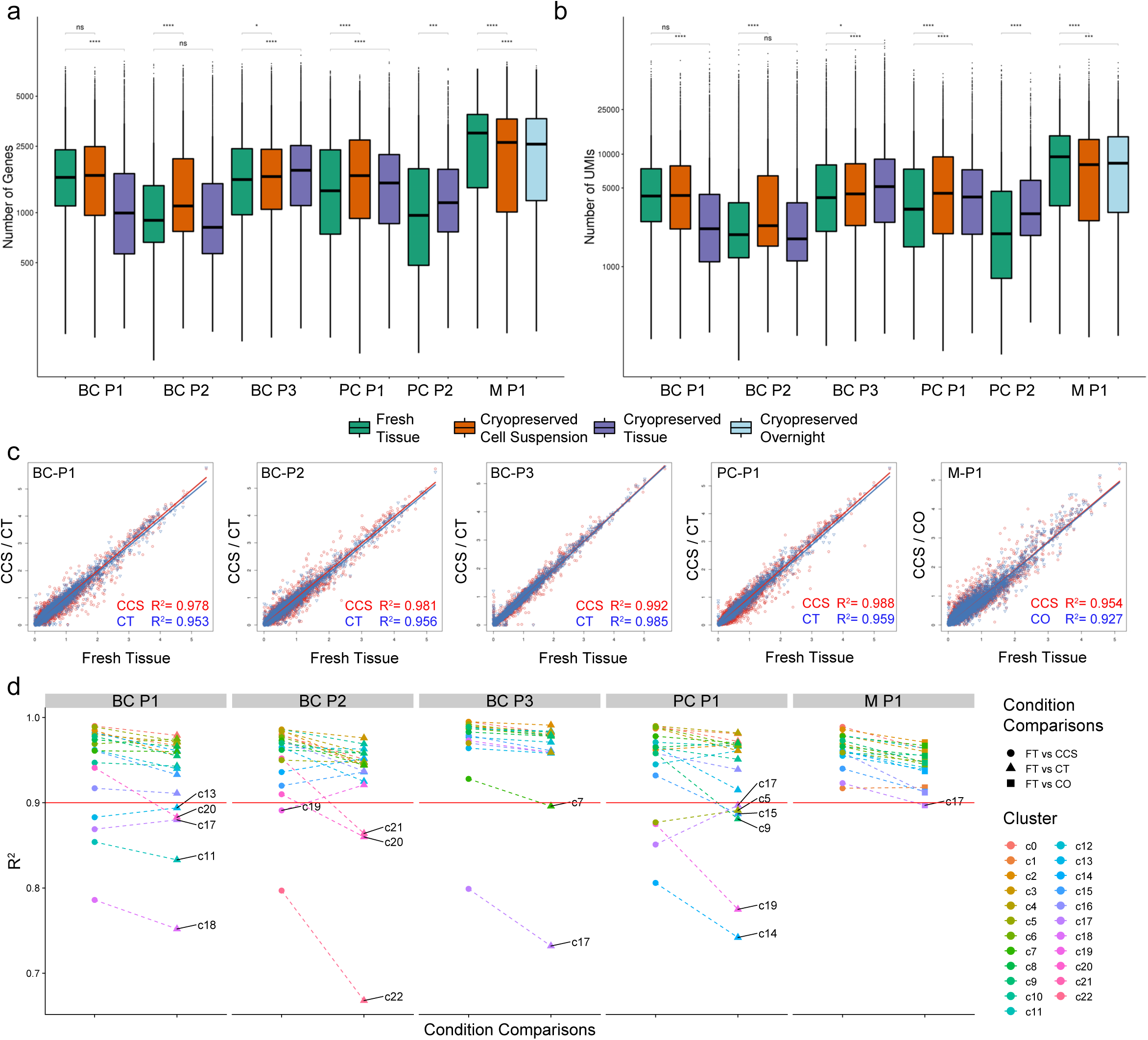
Cryopreservation maintains the integrity and complexity of single-cell transcriptomes in clinical human cancers. **a-b**, Number of genes **(a)** and UMIs **(b)** detected per cell across all FT, CCS, CT, and CO replicates from breast (BC-P1, BC-P2 and BC-P3), prostate (PC-P1 and PC-P2) and melanoma samples (M-P1). Sequencing libraries were down sampled to equal number of mapped reads per cell using cellranger aggregate function to account for differences from sequencing depth. Note that only one CCS replicate in M-P1 (orange) and one CT replicate in BC-P1 (purple) had significantly lower number of genes and UMIs per cell compared to their matching FT replicate. Statistical significance was determined using an unpaired Student’s *t-*test. **c**, Pseudobulk gene correlations between FT cells with CCS (red line) and CT or CO (blue line) replicates. Correlation values (adjusted-*R*^*2*^) were computed using linear regression in *R* to model the log-normalised gene expression values between two replicates. In all cases, CCS replicates had higher *R*^*2*^ values compared to CT and CO comparisons. **d**, Cluster-level gene correlations between FT cells with CCS (circle), CT (triangle) and CO (square) replicates show similar trends with pseudobulk gene correlations. Dotted lines join corresponding clusters between different comparison methods. Note that plasmablasts (c18 in BC-P1 and c22 in BC-P2) was the only cell type identified in multiple cases to have significantly lower correlations.

We next investigated the gene correlation between FT samples and their respective cryopreserved replicates. Bulk gene correlations revealed high *R*^*2*^ values between FT and all cryopreserved replicates (*R*^*2*^ > 0.90; Fig. 3c) where on average, CCS replicates had higher *R*^*2*^ values with the FT sample (mean *R*^*2*^ = 0.98, min = 0.95 and max = 0.99) compared to the CT replicates (mean *R*^*2*^ = 0.96, min = 0.93 and max = 0.99) (Fig. 3c). Similarly, we examined if this trend was unique to particular cell types on the clusters level (Fig. 3d). Only clusters containing cells from all three replicates with a minimum cluster size 100 and at least 20 cells per replicate were examined for representative gene correlations, in order to not be skewed by low cell numbers. Cluster correlations revealed consistent trends with the bulk comparisons, where CCS replicates consistently showed slightly higher *R*^*2*^ correlation than with FT replicates (Fig. 3d). Although a majority of clusters displayed high correlations (*R*^*2*^ > 0.90; indicated by the red line in Fig. 3d), several smaller clusters showed significantly lower correlations than the bulk (*R*^*2*^ < 0.90; Fig. 3d) including five clusters in BC-P1 (c13 - cancer/epithelial, c20 - cancer/epithelial, c17 - T-cells, c11 – endothelial and c18 – plasmablasts), four clusters in BC-P2 (c19 – perivascular, c21 – pDCs, c20 – T-cells and c22 – plasmablasts), two clusters in BC-P3 (c7 – monocyte/macrophage and c17 – unassigned cluster), six clusters in PC-P1 (c17 – NK cells, c5 – cancer/epithelial, c15 – endothelial, c9 perivascular, c19 – mast cells and c14 – cancer/epithelial) and one cluster in M-P1 (c17 – CAFs). The majority of these poorly correlated clusters were comprised of small cell numbers. The only cell type consistently found to have very poor correlation values across multiple cases (*R*^*2*^ < 0.80) was plasmablasts (c18 in BC-P1 and c22 in BC-P2), suggesting that cell type is more prone to transcriptional changes due to cryopreservation (Fig. 3d). Taken together, we find that cryopreservation can conserve high quality transcriptomes for scRNA-Seq. These data suggest that processing scRNA-Seq from CCS yields slightly higher quality data than from CT. Although the sample number was small, we found that cryopreservation induced changes in transcriptome integrity of plasmablasts identified in breast tumours, warranting some caution for studying this cell type using this method.

### Tumour cryopreservation maintains biological pathways

Biological and functional findings from scRNA-Seq experiments are often interpreted through the gene ontology (GO) analysis for pathway enrichments across unique cell clusters. To assess if such downstream analyses are impacted by cryopreservation, we first separated our integrated clusters by their cryopreservation conditions. We then performed differential gene expression and GO pathway enrichment to assess how pathways detected across FT clusters were detected in their respective cryopreserved replicates. This analysis revealed a good overlap of total detected pathways in all cancer cases, with over 64% of all FT pathways consistently detected in both cryopreserved replicates in all cases (min = 64% and max = 77%; Fig. 4a). For pathways that were unique to FT replicates and not detected in the matching cryopreserved replicates, no common pathways were shared across the FT replicates from all six cases, however, a total of seven pathways were shared across three cases. Though this may reflect gene expression programs that might be affected by the cryopreservation process, these pathways were mostly detected across different cell types, with the exception of the gene sets GO:0016628 (‘oxidoreductase activity’) and GO:0016791 (‘phosphatase activity’), which were unique to cancer/epithelial cells and T-cells from three FT replicates, respectively (Supplementary Table 3). From the high concordance of GO pathways detected in cryopreserved replicates, we concluded that these minor differences were likely due to the variations in the scRNA-Seq platform or false discovery rather than the cryopreservation process.

**Figure 4.**
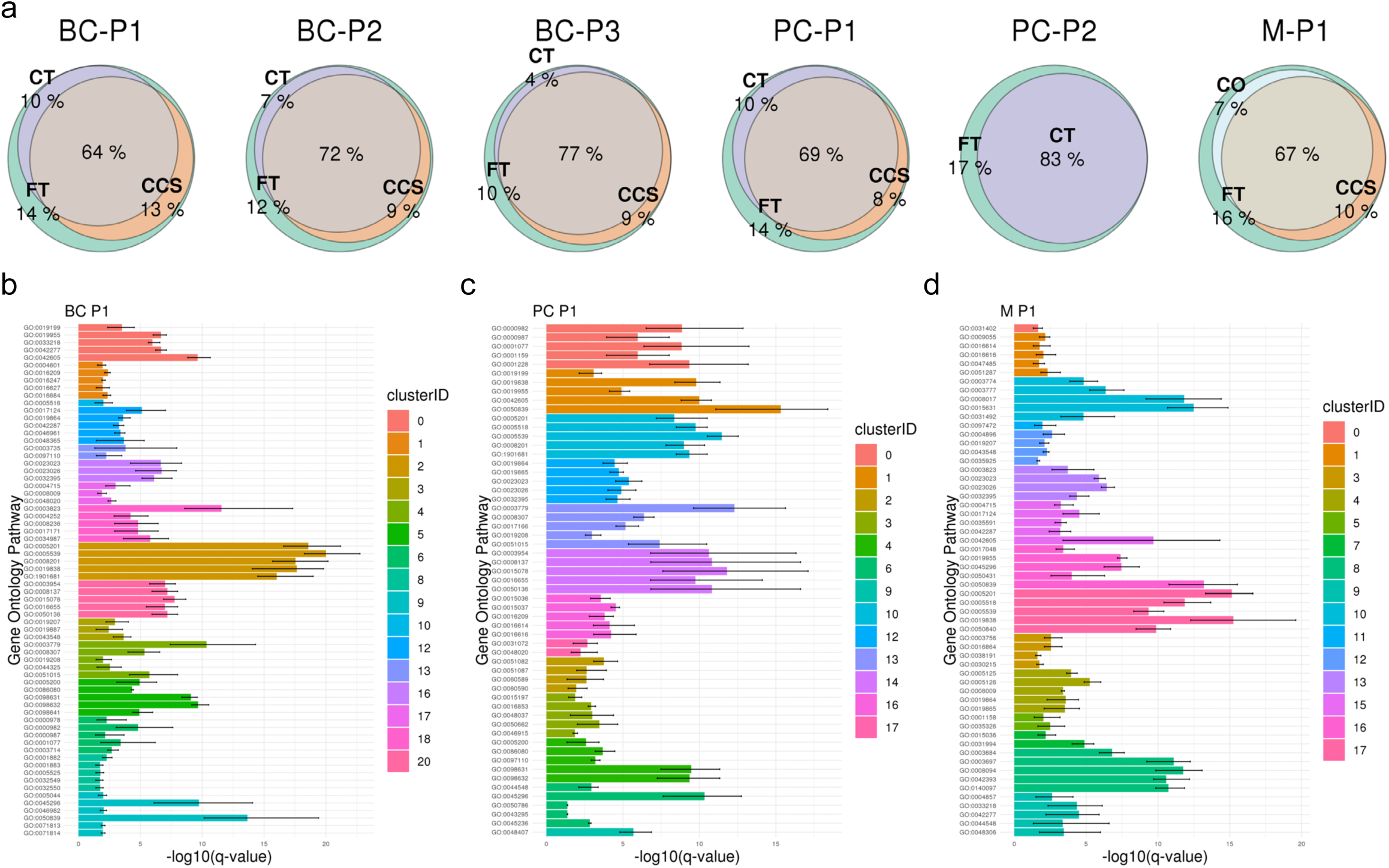
Methods of human tumour cryopreservation maintains biological pathways. **a**, Euler diagrams highlighting the overlaps between gene ontology (GO) pathways detected in FT clusters and cryopreserved replicates from CCS, CT, and CO. A total of 315, 347, 368, 262, 230 and 311 pathways were assessed from the FT replicates across the BC-P1, BC-P2, BC-P3, PC-P1, PC-P2 and M-P1 cases, respectively. **b-d**, Sensitivity of pathway enrichment scores detected in clusters across cryopreserved replicates of BC-P1 (**b**), PC-P1 (**c**) and M-P1 (**d**). The minimum, mean and maximum -log10 q-value are plotted in the error bars of each GO pathway. All DEGs from each cluster were passed on to the ClusterProfiler package for functional enrichment with the CC sub-ontology under the human org.Hs.eg.db database. GO pathway descriptions can be found in Supplementary Table 3.

We next assessed the variability of pathway enrichment scores for cryopreserved cells from each cluster (Fig. 4b-d). This analysis revealed minimal variability across clusters from all six cases of breast cancers, prostate cancers and melanoma, represented by the small range of -log10 q-value enrichment scores for cells across FT and cryopreserved replicates (Fig. 4b-d; Fig. S4). Taken together, these data indicate that the minor variations emerging from cryopreservation, as shown previously through clustering metrics (Fig. 1e-g; Fig. 2f-h), transcript detection (Fig. 3a) and gene correlations (Fig. 3b-c), only have minor impacts on downstream analyses such as the detection of key biological pathways.

### Whole cell cryopreservation allows for highly robust immunophenotyping using CITE-Seq

Immunophenotyping with barcoded-antibody methods such as CITE-Seq can be powerfully applied to simultaneously integrate protein and gene expression in single cells. Although previous studies have applied CITE-Seq to cryopreserved peripheral blood mononuclear cells, it has yet to be established whether CITE-Seq can be applied to cells cryopreserved as solid tissues [5]. As cell surface markers have been extensively used to characterise immune subpopulations, such additional layers of phenotypic information can be used to profile the tumour immune response in cryopreserved patient samples. Here, we performed CITE-Seq of an independent breast cancer case cryopreserved as CT (Fig. 5a) using a panel of 15 canonical cell type markers. We first used a combination of canonical markers from RNA expression to broadly annotated clusters (Fig. 5a; Fig. S5a). From CITE-Seq, we were able to validate our cell type annotations by showing the highly specific antibody-derived tag (ADT) expression levels of canonical markers on corresponding cell types. For example, ADT levels of EPCAM on cancer/epithelial cells (c0, c4, c8, c14 and c15), CD31 (*PECAM1*) and CD34 on endothelial cells (c7 and c9), CD146 (*MCAM*) on perivascular cells (c11), CD90 (*THY1*) and CD34 on CAFs (*c13*) and CD45 (*PTPRC*) on immune cells (c3, c5 and c12) (Fig. 5b-c; Fig. S5a). Within the immune compartments, CD3 specifically marked T-cells, while CD4 and CD8 were more specifically expressed on the respective T-cell subpopulations (Fig. 5b; Fig. S5a). ADT levels of the activation marker CD69 and tissue resident marker CD103 were heterogeneously expressed on T-cell subpopulations (Fig. 5b). CD11c and CD11d were highly specific to monocyte/macrophage cell clusters (Fig. 5b). Major histocompatibility complexes, MHC-II and MHC-I, were highly expressed by endothelial cells, whereas MHC-II was also detected on monocyte/macrophage clusters (Fig. 5b).

**Figure 5.**
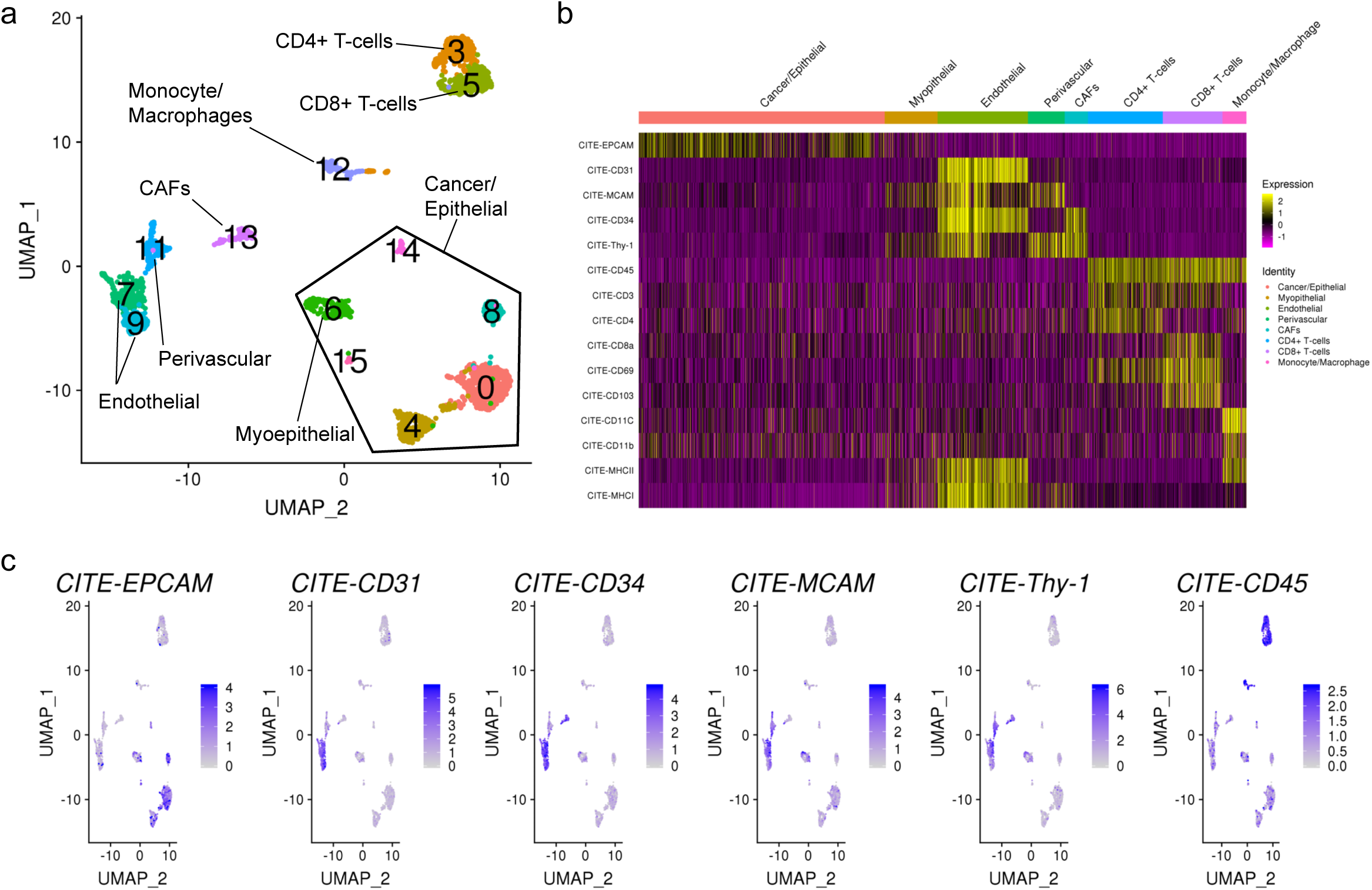
Cryopreservation provides high quality immunophenotyping using CITE-Seq. **a**, UMAP visualisation of 2,621 cells sequenced from an independent breast cancer case cryopreserved as CT. Clusters were annotated based on canonical cell type markers by RNA expression. CITE-Seq was performed on this case using a panel of 15 canonical cell type markers. **b**, Heatmap of rescaled antibody-derived tag (ADT) values for relevant markers for cancer/epithelial cells (EPCAM), endothelial cells (CD31/*PECAM1* and CD34), perivascular cells (*MCAM*/CD146 and THY-1/CD90), cancer-associated fibroblasts (THY-1/CD90 and CD34), immune cells (CD45/*PTPRC*), T-cells (CD3, CD4, CD8, CD69 and CD103), monocytes/macrophages (CD11c and CD11d) and MHC molecules (MHC-II and MHC-I). **c**, Featureplot representation of ADT protein expression values for selected markers from **(b)** highlighting the specificity major lineage markers on RNA based clustering in **(a)**.

ADT levels, which overcomes several technical limitations from gene drop-out, have a greater sensitivity than UMI counts by scRNA-Seq. The average correlation between ADT levels and the corresponding gene expression for this panel of 15 markers was 0.214 (min *R*^*2*^ = 0.003 and max *R*^*2*^ = 0.639; Fig. S5b). This ranged significantly for different markers, particularly for lowly expressed immunoregulatory molecules such as CD4 (*CD4*), CD103 (*ITGAE*), CD11b (*ITGAD*) and CD11c (*ITGAX*), where expression levels of their corresponding genes were lowly detected in comparison to the ADT, with *R*^*2*^ values of 0.016, 0.005, 0.003 and 0.004, respectively (Fig. S5b). In contrast, highly expressed genes such as the endothelial cell marker CD31 (*PECAM1*) showed much higher correlations (*R*^*2*^ = 0.639; Fig. S5b). In summary, we show that good quality CITE-Seq data can be generated from cells cryopreserved as solid CT. Such methods can be used to powerfully extract additional phenotypic information from low amounts of cryopreserved clinical tissue, aiding the annotation of single-cell clusters and the detection of clinically relevant molecules such as immune-checkpoints.

## Conclusions

We show that high quality scRNA-Seq data can be generated from human cancer samples cryopreserved as dissociated single-cell suspensions (CCS) and solid tissues (CT). For the latter, minimal processing is required following sample collection and can be conducted routinely in hospital pathology laboratories that have access to -80°C freezers for short-term storage. These samples can later be transported to a research laboratories for long-term storage or further processing. We found that CCS samples yielded slightly higher quality data, however CCS requires more specialised tissue processing following sample collection before cryopreservation (∼1-2 hours using commercial dissociation kits). While we used tissues that had been cryopreserved for up to 6 weeks in this study, we have routinely processed samples stored at liquid nitrogen for more than 3 years for scRNA-Seq. Most importantly, we show that the complexity of the TME is conserved following cryopreservation as both CCS and CT. This is an important consideration because an integrated understanding of the neoplastic, stromal and immune states define tumours and their response to treatment. Further, we show that multi-omics methods, such as immunophenotyping using CITE-Seq, can be performed using cells cryopreserved as solid tissue pieces, which is impossible when using other preservation methods such as single nuclei sequencing from snap frozen tissues. Our findings have allowed sample multiplexing methods to be applied to clinical samples to reduce cost and logistics for project scaling, such as barcode hashing or genotype based demultiplexing (unpublished data) [9, 10]. Due to the easily adoptable nature of cryopreserving solid tissues in tissue biobanking processes, we envisage our findings to positively impact the sample collection opportunities for future clinical studies, particularly for multi-site collaborative studies, to allow for the centralisation of sample processing and batched analysis.

## Methods

### Ethics approval and consent for publication

Patient tissues used in this work were collected under protocols x12-0231, x13-0133, x16-018, x17-0312 and x17-155. Human Research Ethics Committee approval was obtained through the Sydney Local Health District Ethics Committee, Royal Prince Alfred Hospital zone, and the St Vincent’s Hospital Ethics Committee. Written consent was obtained from all patients prior to collection of tissue and clinical data stored in a de-identified manner, following pre-approved protocols. Consent into the study included the agreement to the use of all patient tissue and data for publication.

### Primary tissue dissociation and sample preparation

Fresh surgically resected tissues were washed with RPMI 1640 (ThermoFisher Scientific) and diced into 1-2 mm^3^ pieces. Tissue pieces were mixed and approximately one third were viably frozen in cryogenic vials in 5% Dimethyl sulfoxide (DMSO) and 95% Fetal Bovine Serum (FBS) at 1°C/minute in -80°C using Mr. Frosty™ Freezing Containers (ThermoFisher). This was classified as the solid cryopreserved tissue (CT) sample. The remaining tissue was further minced with scissors and enzymatically dissociated using the Human Tumour Dissociation Kit (Miltenyi Biotec) following the manufacturer’s protocol. Following incubation with the enzymes, the sample was resuspended in media (80% RPMI 1640, 20% FBS) and filtered through MACS® SmartStrainers (70 µM; Miltenyi Biotec). The resulting single cell suspension was centrifuged at 300 × g for 5 min. At this stage, a proportion of the dissociated cell suspension was frozen in cryogenic vials in 10% DMSO, 50% FBS and 40% RPMI 1640 at 1°C/minute in -80°C using Mr. Frosty™ Freezing Containers (ThermoFisher). This was classified as the dissociated cryopreserved cell suspension (CCS) sample. For the dissociated fresh tissue (FT) sample, red blood cells were lysed with Lysing Buffer (Becton Dickinson) for 5 min and neutralised with media (80% RPMI 1640, 20% FBS). Cells were further filtered through a 40 μm filter and centrifuged at 300 × g for 5 min. Viability was assessed using Trypan Blue (ThermoFisher). For samples with a viability score of < 80%, enrichment was performed using the EasySep Dead Cell Removal (Annexin V) Kit (StemCell Technologies) following the manufacturer’s protocol. Enriched cell suspensions were resuspended in a final solution of PBS with 10% FBS solution prior to loading on the 10X Chromium platform. For the processing of cryopreserved replicates, samples were frozen at -80° C for ∼1 week followed by ∼5 weeks at -196°C for prior to scRNA-Seq. For obvious logistical reasons (freezing storage time), FT samples were run on the 10X Chromium platform immediately whilst CT and CSS samples were processed simultaneously at a later date. Following cryopreservation, samples were thawed in a 37°C water bath and washed multiple times with RPMI 1640. CT samples were dissociated in the same manner as the FT samples, as previously described. CCS samples were enriched for live cells if viability was assessed to be < 80%, as described above. For both cryopreserved replicates from breast tumours, the mouse cell line NIH3T3 was thawed and spiked in at 2% of the total cell number prior to cell loading on the 10X Chromium.

### Single-cell RNA sequencing on the 10X Chromium platform

High throughput scRNA-Seq was performed using the Chromium Single Cell 3’ v2 and 5’ chemistry (10X Genomics) according the to the manufacturer’s instructions. All replicates within a case were captured using the same chemistry. A total of 6,000 cells were targeted per lane. SCRS libraries were sequenced on the Illumina NextSeq 500 platform with pair-end sequencing and dual indexing according to the recommended Chromium platform protocol; 26 cycles for Read 1, 8 cycles for i7 index and 98 cycles for Read 2.

### Data processing

Read demultiplexing and alignment to the GRCh38 human reference genome was performed using the Cell Ranger Single Cell Software v2.0 (10X Genomics) with the cellranger mkfastq and count functions, respectively. For cryopreserved replicates from breast tumours with mouse cell line spike in (NIH3T3), the above steps were performed using the GRCh38 human and mm10 mouse reference genomes. Raw count matrices were filtered for ‘real’ barcodes using the EmptyDrops package in *R* which calculates deviations against a generated ambient background RNA profile [11]. Additional conservative cut offs were further applied based on the number of genes detected per cell (greater than 200) and the percentage of mitochondrial unique molecular identifier (UMI) counts (less than 20%). Filtered barcodes from matched replicates were then processed and integrated using the Seurat v3 package in *R* as per the developers’ vignettes [8]. For the comparison of transcript metrics across cryopreserved replicates, including the number of genes, UMIs and gene correlations, we performed downsampling of sequencing libraries by the total number of mapped reads using the cellranger aggr function. For comparison of clusters across cryopreservation conditions, cells were randomly down sampled to the lowest replicate size using the data.table package in *R*.

### Silhouette scores, mixing metric and local structure metric

We applied clustering and mixability metrics from Stuart *et al.* to quantitative measure the robustness of the cryopreserved replicates to reflect good technical replicates with the FT [8]. Stratified random down sampling was first applied to each case to generate clusters with equal sizes across all three conditions. This was performed using data.table package in *R*. As a positive control, FT datasets were randomly down sampled to generate two pseudo-replicates. Three comparisons were computed per case: FT-1 vs FT-2, FT-1 vs CCS and FT-1 vs CT. For the melanoma case, the comparisons were FT-1 vs FT-2, FT-1 vs CCS and FT-1 vs CO. Silhouette scores, mixing metrics and local structure metrics were all computed using code adopted from the Seurat v3 package [8].

### Bulk and cluster level gene correlations

Adjusted *R*^*2*^ correlation values were calculated using linear regression, implemented in *R*. Sequencing libraries normalised by the number of mapped reads using CellRanger were used. Pseudo-replicate bulks and cluster-level bulks were generated from log-normalised gene expression values. FT bulk and cluster level replicates were compared to cryopreserved replicates (CCS/CT/CO).

### Differential gene expression and pathway enrichment

Integrated cases were split by replicate. Differential gene expression was then performed between integrated cluster IDs across each of the replicates using the MAST method through the *FindAllMarkers* function in Seurat (log fold change threshold of 0.25, *p-value* threshold of 1×10^−5^ and FDR threshold of 0.05) [12]. All DEGs from each cluster were then passed on to the ClusterProfiler package for functional enrichment [13]. The *compareCluster* function was used with the enrichGO default CC sub-ontology under the human org.Hs.eg.db database. The overlaps of detected GO pathways across each replicate were computed and visualised using the euler and ggplot2 packages in R.

### CITE-Seq staining and data processing

Samples were stained with 10X Chromium 3’ mRNA capture compatible TotalSeq-A antibodies (Biolegend, USA). Staining was performed as previously described by Stoeckius et. al (2017) with a few modifications [5]. Briefly, a maximum of 2 million cells per sample was resuspended in 100 µl of cell staining buffer (Biolegend, USA) with 5 µl of Fc receptor Block (TrueStain FcX, Bioelegend, USA) for 15 minutes followed by a 30min staining of the antibodies at 4°C. A concentration of 1 µg / 100 µl was used for all antibody markers used in this study. The cells were then washed 3x with PBS containing 10% FBS media followed by centrifugation (300 x g for 5 min at 4°C) and expungement of supernatant. The sample was then resuspended in PBS with 10% FBS for 10X Chromium capture. Indexed CITESeq libraries were spiked in to 10X scRNA-Seq libraries for sequencing on the NextSeq500 platform (Illumina). Reads were demultiplexed using CellRanger v2.0. Cell counts of CITE antibodies were calculated from sequenced CITE libraries with CITE-seq-Count v.1.4.3 using default parameters recommended by developers. Counts were integrated with scRNA-seq data using Seurat (v.3.1.4), scaled and normalised.

## Data availability

The scRNA-Seq data from this study has been deposited in the European Nucleotide Archive (ENA) under the accession code PRJEB38487. This depository demultiplexed paired ended reads (R1 and R2), Illumina indices and bam files processed using the Cellranger software. Code related to the scRNA-Seq analysis can be found on the website: https://github.com/sunnyzwu/cryopreservation_scRNAseq. All other relevant data are available from the authors upon request.

## Acknowledgements

This work was supported by funding from John and Deborah McMurtrie, the National Breast Cancer Foundation (NBCF) of Australia; and The Sydney Breast Cancer Foundation. A.S. is the recipient of a Senior Research Fellowship from the National Health and Medical Research Council of Australia. S.Z.W. is supported by the Australian Government Research Training Program Scholarship. S.O.T. is supported by the NBCF (PRAC 16-006), the IIRS 19 084 and the Sydney Breast Cancer Foundation and the Family and Friends of Michael O’Sullivan. L.H was supported by the Funding sources - APCRC-NSW (DoHA) and CINSW TPG. R.A.S. was supported by an National Health and Medical Research Council of Australia (NHMRC) Program Grant and NHMRC Practitioner Fellowship grant. Support from Melanoma Institute Australia and The Ainsworth Foundation is also gratefully acknowledged. We would like to thanks the following people for their assistance in the experimental part of this manuscript; Ms. Gillian Lehrbach from the Garvan Tissue Culture Facility; Anne-Maree Haynes and Daniela Barrato from the Prostate Cancer Biobank; The Garvan-Weizmann Centre for Cellular Genomics for their expertise in single-cell sequencing, and Mr. Dominik Kaczorowski for his help in next-generation sequencing. This manuscript was edited at Life Science Editors.

## Author Contributions

A.S. conceived the project and directed the study. S.Z.W. and A.S. wrote the manuscript. All authors reviewed the drafting of the manuscript. E.L., S.O.T., M.N.H., E.A.M., J.B., D.S., C.M. and S.W. organised the access to breast cancer patient tissue. L.H., A.M.J. and P.S. organised the access to patient tissue from prostate cancer patients. G.V.L., R.A.S., J.S. and C.Q. organised the access to melanoma tissue. K.H. collected clinical samples. S.Z.W., K.H. and A.C. performed tumour dissociation for scRNA-Seq. C.C. helped perform the next-generation sequencing of the scRNA-Seq libraries. S.Z.W. performed and interpreted the pre-processing and downstream analysis of the scRNA-Seq data. D.R. supervised the scRNA-Seq analysis. G.A. performed the CITE-Seq staining experiments. B.Z.Y. provided guidance on CITE-Seq experimental parameters and sequencing. N.B. performed bioinformatics analysis of the CITE-Seq experiments. E.L., S.J. and J.P. provided intellectual input.

## Conflicts of interests

No conflicts of interests. R.A.S. has received fees for professional services from Merck Sharp & Dohme, GlaxoSmithKline Australia, Bristol-Myers Squibb, Novartis, Myriad, NeraCare and Amgen. G.V.L. is consultant advisor to Aduro, BMS, Mass-Array, Merck MSD, Novartis, Pierre-Fabre, Roche, Sandoz, and Qbiotics.

## Supplementary Figure Legends

**Supplementary Figure 1.**
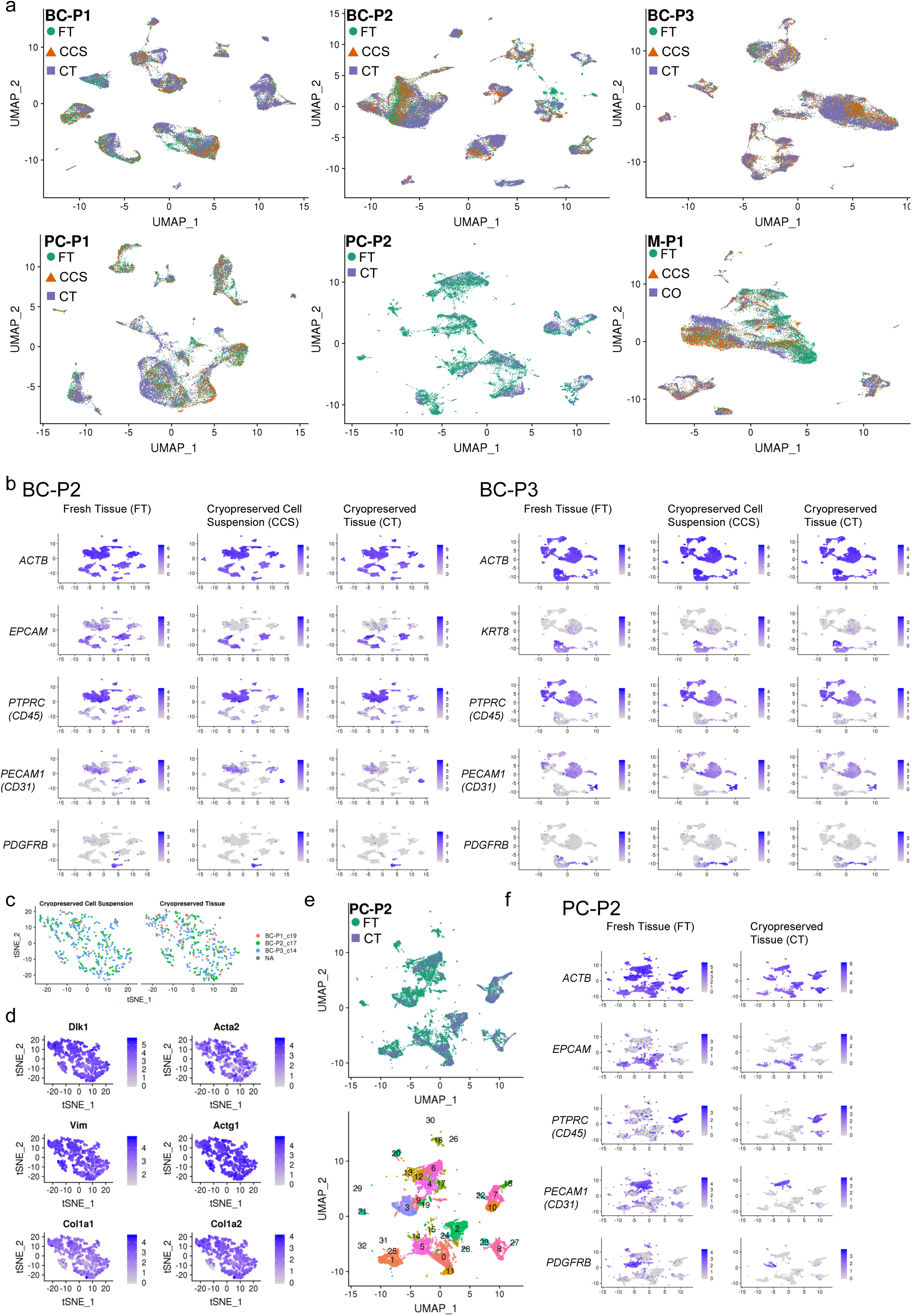
Cryopreservation allows for robust cell-type detection in clinical cancer samples. **a**, UMAP visualisations for the non-batch corrected data for each of the three breast cancer (BC-P1, BC-P2 and BC-P3), two prostate cancer (PC-P1 and PC-P2) and metastatic melanoma case (M-P1). **b**, Featureplot visualisations for additional breast cancer cases BC-P2 and BC-P3. Gene expression shows the conservation of the housekeeping gene *ACTB*, and markers for cancer/epithelial (*EPCAM*), immune (*PTPRC/*CD45), endothelial (*PECAM1*/CD31) and fibroblast/perivascular (*PDGFRB*) clusters following cryopreservation as CCS and CT. **c**, *t*SNE visualisation showing the high mixability of mouse NIH3T3 fibroblast cell line spike ins (∼2%) from the cryopreserved replicates from all three breast cancer cases. Embeddings are split by cells captured from CCS and CT, respectively. Original cluster IDs from Figure 1b are c19 from BC-P1, c17 from BC-P2 and c14 from BC-P3. **d**, Featureplot visualisations of the NIH3T3 cell line fibroblast markers *Dlk1, Acta2, Vim, Actg1, Col1a1* and *Col1a2*. **e**, UMAP visualisations for the batch corrected data for PC-P2, which only contains comparisons between FT and CT replicates due to low cell numbers in the CCS replicate. UMAPs are coloured by cryopreserved conditions and cluster IDs. **f**, Featureplot visualisations of gene expression highlighting the conservation of the major cell lineages, as represented in **(b).**

**Supplementary Figure 2.**
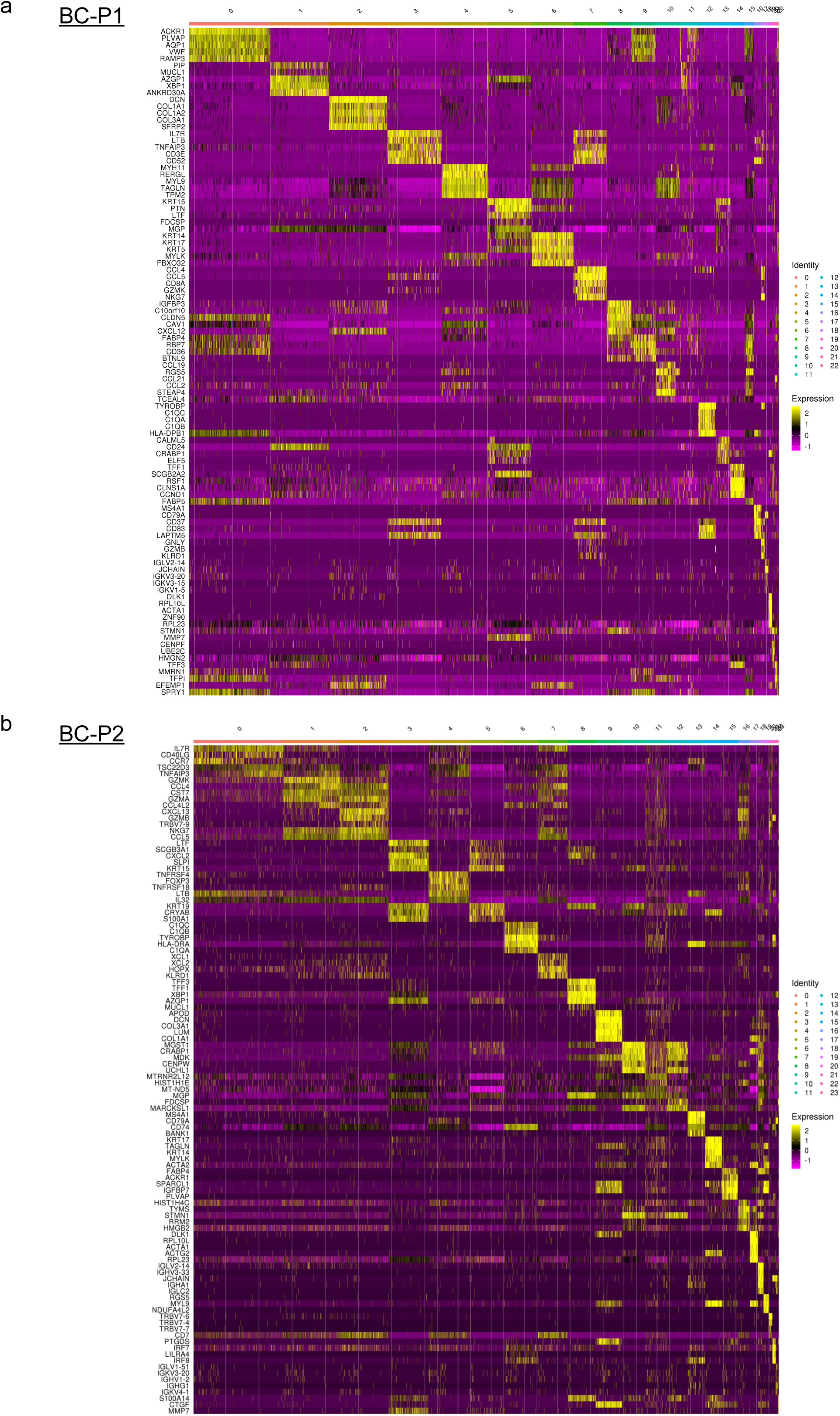
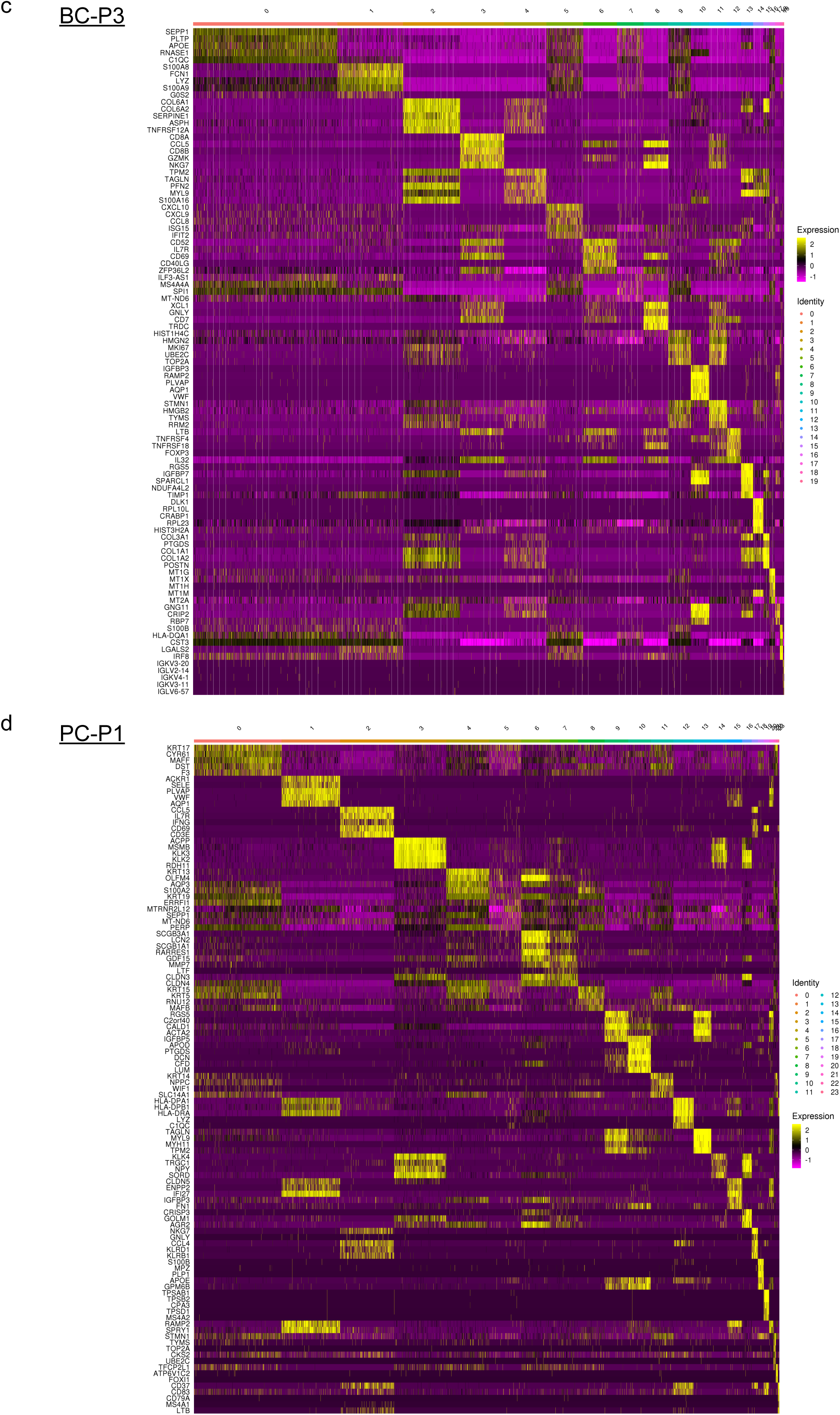
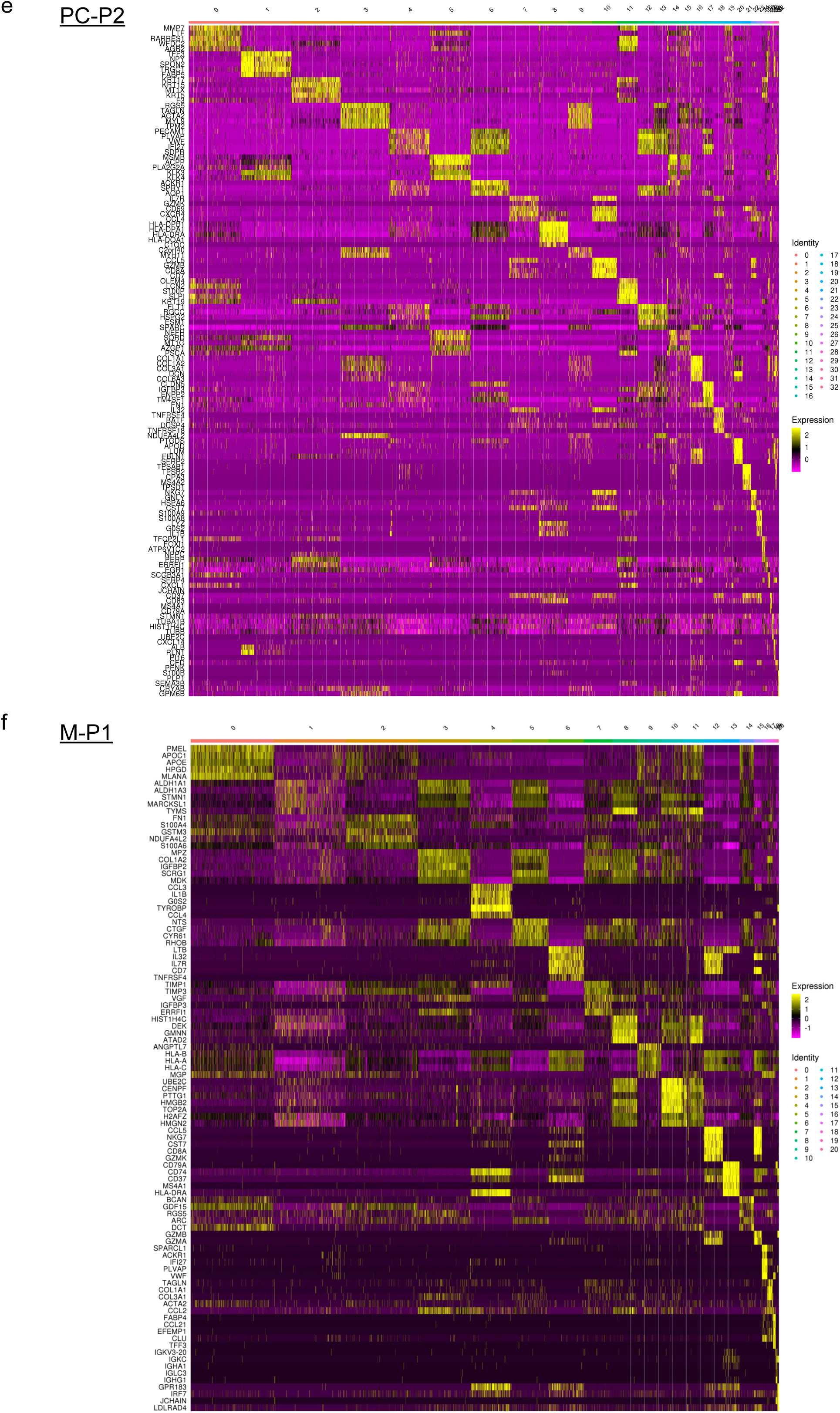
Heatmaps of integrated clusters for breast, prostate and melanoma cancer case. **a-f**, Heatmap visualisation of the top 5 differentially expressed genes per cluster for three breast cancer cases BC-P1 **(a)**, BC-P2 **(b)** and BC-P3 **(c)**, two prostate cancer cases PC-P1 **(d)** and PC-P2 **(e)** and a metastatic melanoma M-P1 (**f**). All cases represent the integrated clustering of all cryopreserved conditions. Differentially gene expression was performed using the MAST method within Seurat v3 with the RNA assay and default parameters. Heatmaps were generated using the DoHeatMap function using Seurat v3. Complete gene lists used are detailed in Supplementary Table 2.

**Supplementary Figure 3.**
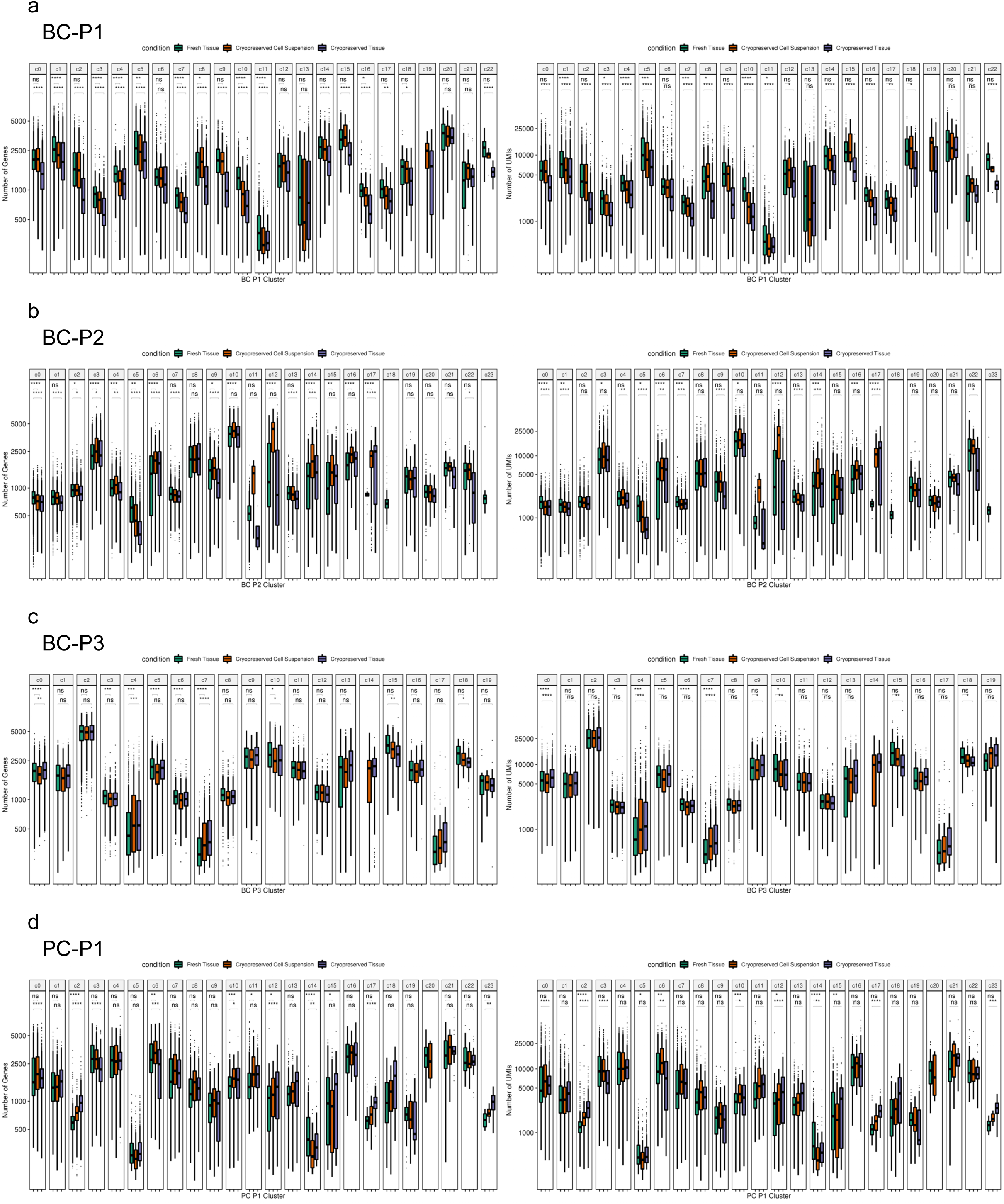
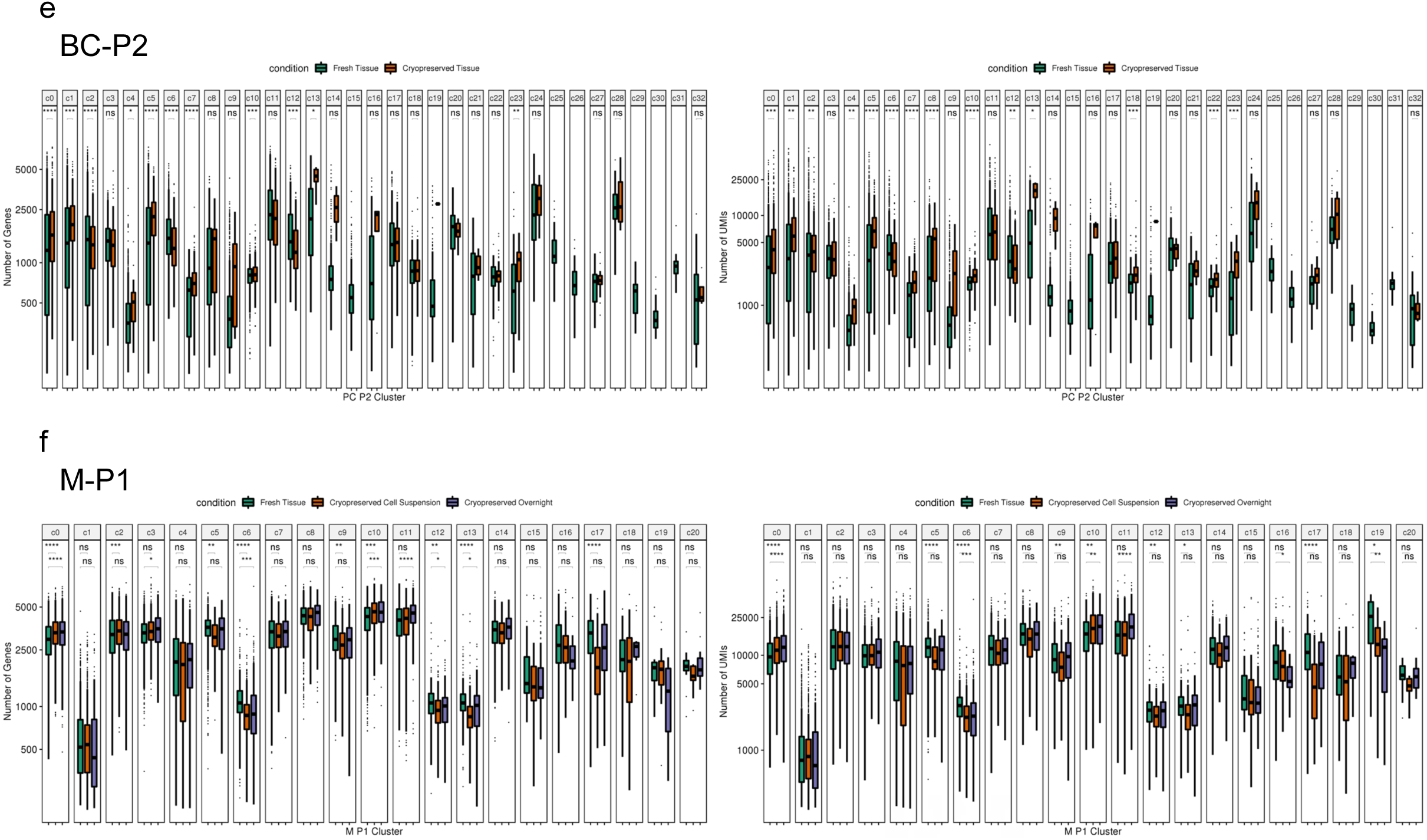
Number of genes and UMIs per cluster. **a-f**, Number of genes (left) and UMIs (right) detected per cell per cluster across FT (green), CCS (orange), CT (purple) and CO (purple; melanoma case only) replicates of breast cancer cases BC-P1 **(a)**, BC-P2 **(b)** and BC-P3 **(c)**, prostate cancer cases PC-P1 **(d)** and PC-P2 (**e**) and a metastatic melanoma M-P1 (**f**). Sequencing libraries were down sampled to equal number of mapped reads per cell using cellranger aggregate function to account for differences from sequencing depth. Statistical significance was determined using an unpaired Student’s *t-* test.

**Supplementary Figure 4.**
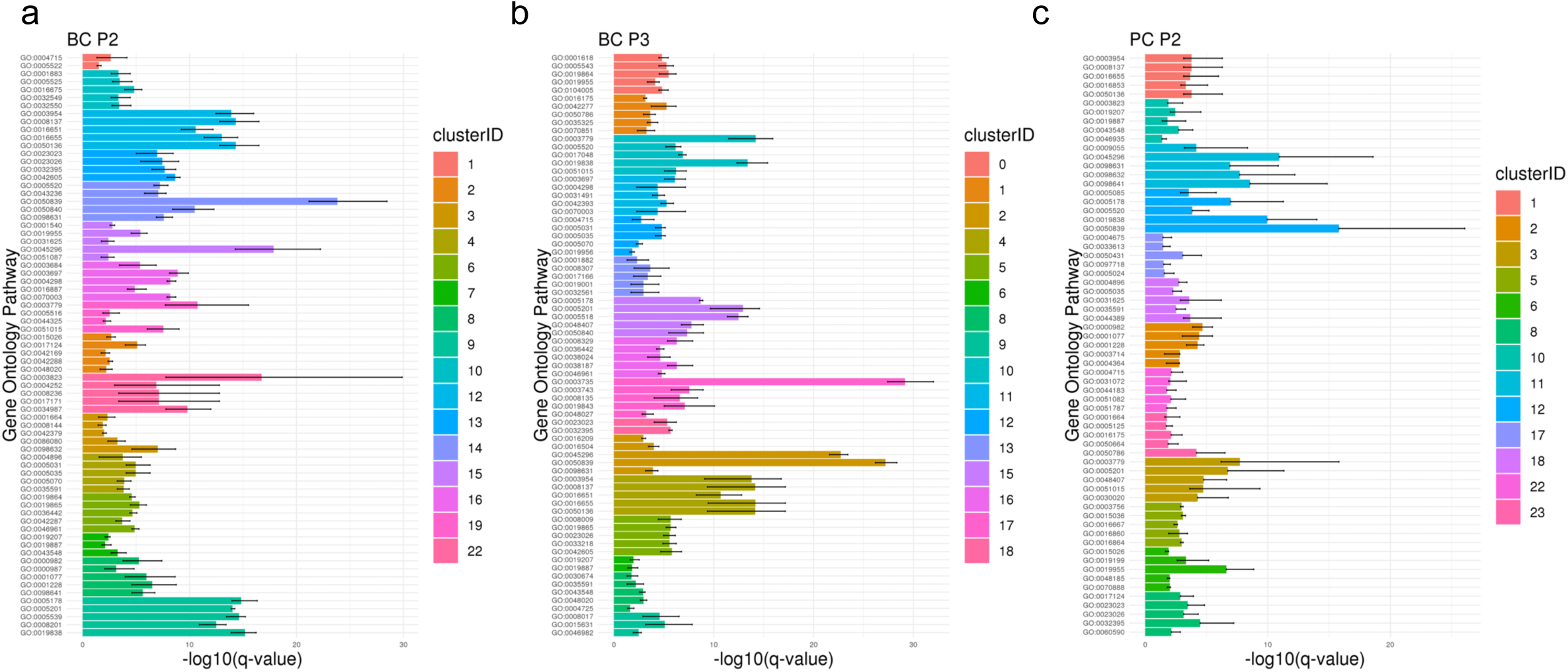
Cryopreservation maintains the detection of biological pathways in additional cases. **a-c**, Sensitivity of pathway enrichment scores detected in clusters across cryopreserved replicates. Additional representative cases of breast cancer BC-P2 **(a)** and BC-P3 (**b**) and prostate cancer PC-P2 (**c**) are shown. The minimum, mean and maximum -log10 q-value are plotted in the error bars of each GO pathway. All DEGs from each cluster were passed on to the ClusterProfiler package for functional enrichment with the CC sub-ontology under the human org.Hs.eg.db database. GO pathway descriptions can be found in Supplementary Table 3.

**Supplementary Figure 5.**
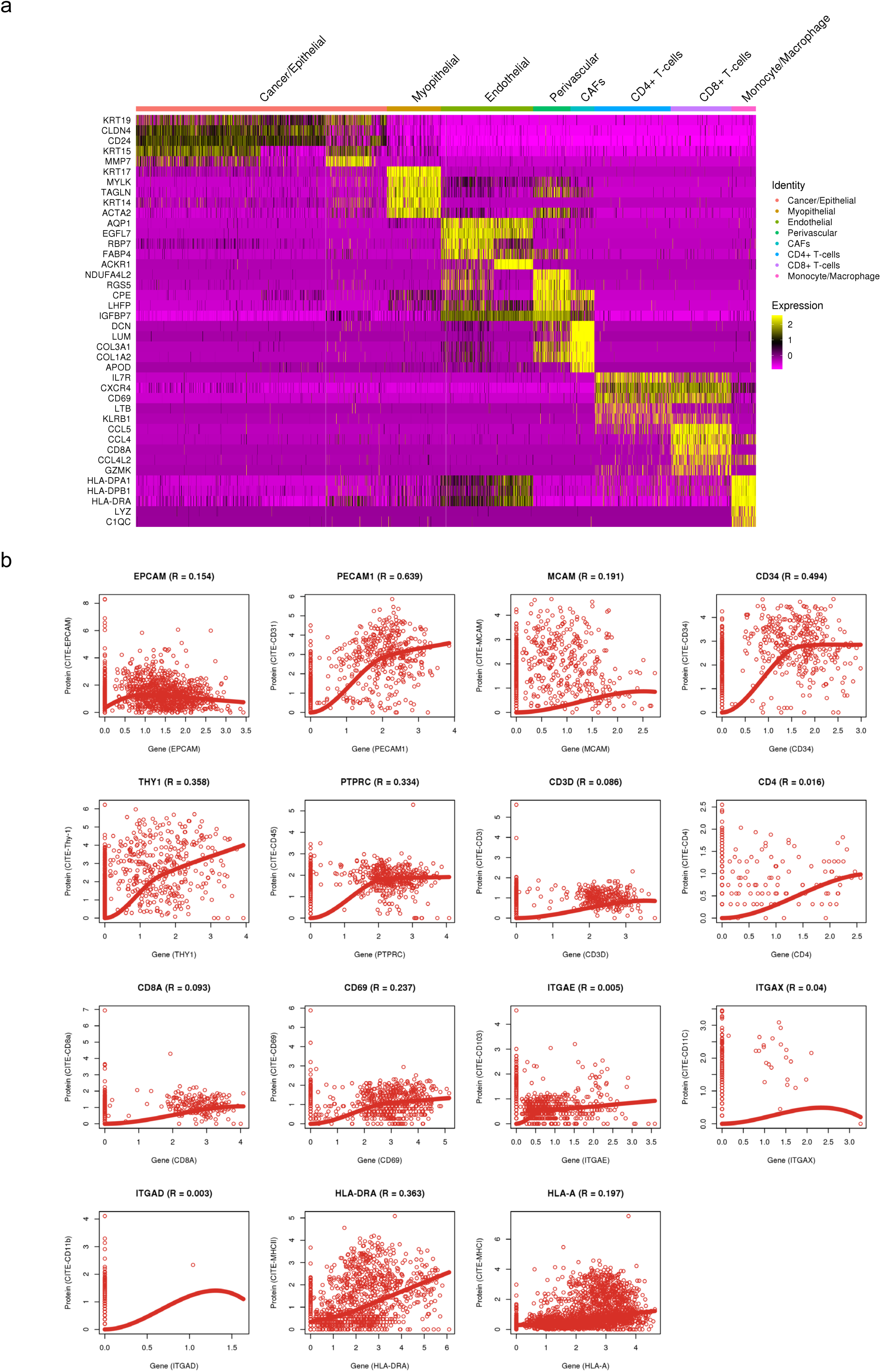
Cryopreservation provides high quality immunophenotyping using CITE-Seq. **a**, Heatmap visualisation of the top 5 differentially expressed genes of indicated canonical cell type markers for an independent breast cancer case for the CITE-Seq experiment. Differentially gene expression was performed using the MAST method within Seurat v3 with the RNA assay and default parameters. Heatmaps were generated using the DoHeatMap function using Seurat v3. **b**, Correlation plots between protein and genes for the panel of 15 markers used for CITE-Seq. Correlation values (adjusted-*R*^*2*^) were computed using linear regression in *R* to model the log-normalised gene expression value and corresponding ADT levels.

## Supplementary Table Legends

**Supplementary Table 1. Clinical information for breast cancer, prostate cancer and metastatic melanoma cases used in this study.**

**Supplementary Table 2. Differentially expressed genes for integrated clusters.** Differentially gene expression was performed using the MAST method within Seurat v3 with the RNA assay and default parameters.

**Supplementary Table 3. Cluster metric standard deviations, cluster level gene correlations and gene pathways unique to cryopreserved conditions. a**, Standard deviations for silhouette scores, mixing metrics and local structure metrics, computed for the comparisons between the down sampled FT cells with FT cells (positive control), CCS, CT and CO. **b**, Cluster level correlation values. Adjusted-*R*^*2*^ values computed using linear regression in R to model log-normalised gene expression values between integrated clustered cells from different cryopreserved replicates. **c**, Functional enrichment between cryopreservation conditions. All DEGs from each cluster were passed on to the ClusterProfiler package for functional enrichment with the CC sub-ontology under the human org.Hs.eg.db database.

## Notes

### Competing Interest Statement

The authors have declared no competing interest.

## References

1. Hanahan, D. and L.M. Coussens, Accessories to the crime: functions of cells recruited to the tumor microenvironment. Cancer Cell, 2012. 21(3): p. 309–22.

2. Madissoon, E., et al., scRNA-seq assessment of the human lung, spleen, and esophagus tissue stability after cold preservation. Genome Biol, 2019. 21(1): p. 1.

3. Attar, M., et al., A practical solution for preserving single cells for RNA sequencing. Sci Rep, 2018. 8(1): p. 2151.

4. Habib, N., et al., Massively parallel single-nucleus RNA-seq with DroNc-seq. Nat Methods, 2017. 14(10): p. 955–958.

5. Stoeckius, M., et al., Simultaneous epitope and transcriptome measurement in single cells. Nat Methods, 2017. 14(9): p. 865–868.

6. Guillaumet-Adkins, A., et al., Single-cell transcriptome conservation in cryopreserved cells and tissues. Genome Biol, 2017. 18(1): p. 45.

7. Picelli, S., et al., Full-length RNA-seq from single cells using Smart-seq2. Nat Protoc, 2014. 9(1): p. 171–81.

8. Stuart, T., et al., Comprehensive Integration of Single-Cell Data. Cell, 2019. 177(7): p. 1888–1902 e21.

9. Stoeckius, M., et al., Cell Hashing with barcoded antibodies enables multiplexing and doublet detection for single cell genomics. Genome Biol, 2018. 19(1): p. 224.

10. Kang, H.M., et al., Multiplexed droplet single-cell RNA-sequencing using natural genetic variation. Nat Biotechnol, 2018. 36(1): p. 89–94.

11. Lun, A.T.L., et al., EmptyDrops: distinguishing cells from empty droplets in droplet-based single-cell RNA sequencing data. Genome Biol, 2019. 20(1): p. 63.

12. Finak, G., et al., MAST: a flexible statistical framework for assessing transcriptional changes and characterizing heterogeneity in single-cell RNA sequencing data. Genome Biol, 2015. 16: p. 278.

13. Yu, G., et al., clusterProfiler: an R package for comparing biological themes among gene clusters. OMICS, 2012. 16(5): p. 284–7.

